# A high-resolution map of bacteriophage øX174 transcription

**DOI:** 10.1101/2020.03.05.979765

**Authors:** Dominic Y. Logel, Paul R. Jaschke

## Abstract

Bacteriophage øX174 is a model virus for studies across the fields of structural biology, genetics, gut microbiome, and synthetic biology, but did not have a high-resolution transcriptome until this work. In this study we used next-generation sequencing to measure the RNA produced from øX174 while infecting its host *E. coli C.* We broadly confirm the past transcriptome model while revealing several interesting deviations from previous knowledge. Additionally, we measure the strength of canonical øX174 promoters and terminators and discover both a putative new promoter that may be activated by heat shock sigma factors, as well as rediscover a controversial Rho-dependent terminator. We also provide evidence for the first antisense transcription observed in the *Microviridae* family, identify two promoters that may be involved in generating this transcriptional activity, and discuss possible reasons why this RNA may be produced.

## INTRODUCTION

Bacteriophage øX174 is a member of the *Microviridae* family and is an important part of the human gut microbiome (Michel et al., 2010; Reyes et al., 2012). Phage ϕX174 produces a tailless icosahedral capsid (Fane et al., 2005) and is known to infect the *Escherichia coli* C strain through the attachment of its major spike protein vertex to the bacterium’s lipopolysaccharide followed by dissociation of the spike protein and conformational changes in the major capsid protein (Hayashi et al., 1988; Sun et al., 2017). The circular single-stranded positive (+) sense DNA genome is transferred across both membranes and the periplasm through an extensible conduit formed by the pilot protein (Sun et al., 2014).

Bacteriophage ϕX174 has been a useful model for genetics research for over 50 years (Benbow et al., 1971; Hutchison, 1969; Jaschke et al., 2019). The 5,386 nucleotide (nt) genome was the first DNA genome sequenced (Sanger et al., 1977) and the number of annotated protein-encoding genes in the ϕX174 genome has changed over time in response to increasingly sophisticated technologies. Specifically, prior to sequencing, ϕX174 was believed to encode seven genes (Hutchison, 1969), however this was revised upwards several times in response to additional experimental evidence (Benbow et al., 1971; Jeng et al., 1970; Mayol and Sinsheimer, 1970). One main feature of the ϕX174 genome that puzzled and fascinated phage researchers is the presence of extensive gene overlaps. Before the genome was sequenced, it was estimated that more sequence length than is covered by the genome was required to encode the observed expressed proteins (Barrell et al., 1976; Benbow et al., 1972). Sequencing solved this mystery by showing that 8 of the 11 genes in ϕX174 overlap, thereby increasing the available genetic encoding capacity beyond the 5,386 nt genome limit (Barrell et al., 1976; Sanger et al., 1977). All ϕX174 genes were discovered through forward genetic screens with the exception of gene K, which was discovered using comparative genomics (Tessman et al., 1980).

The genes in ϕX174, and other *Bullavirinae* sub-family (Godson et al., 1978; Kodaira et al., 1992; Lau and Spencer, 1985), are arranged in functional gene clusters, with four broad types of proteins produced: scaffolding, viral propagation, capsid, and host interaction (Table 1). The functional groups are clustered into two major genomic regions within the *Bullavirinae* sub-family, the first with no overlapping genes and containing the three capsid proteins: F, G, and H. The second genomic region encoding the remaining 8 genes where the majority of the genes overlap with at least one neighbouring gene (Fig. 1A).

**Table 1.** Genes of ΦX174.

| Gene | Gene Product Function | Category |
| --- | --- | --- |
| A | DNA replication, stages II and III | Viral propagation |
| A* | Inhibiting host DNA replication and capsid packaging | Host interaction |
| B | Internal procapsid scaffolding | Scaffolding |
| C | DNA replication stage II to III switching | Viral propagation |
| D | External procapsid scaffolding | Scaffolding |
| E | Host cell lysis | Host interaction |
| F | Major coat protein | Capsid |
| G | Major spike protein | Capsid |
| H | DNA pilot protein, forms a tail for genome transport | Capsid |
| J | DNA packaging | Viral propagation |
| K | Burst size modulation | Viral propagation |

**Fig. 1.**
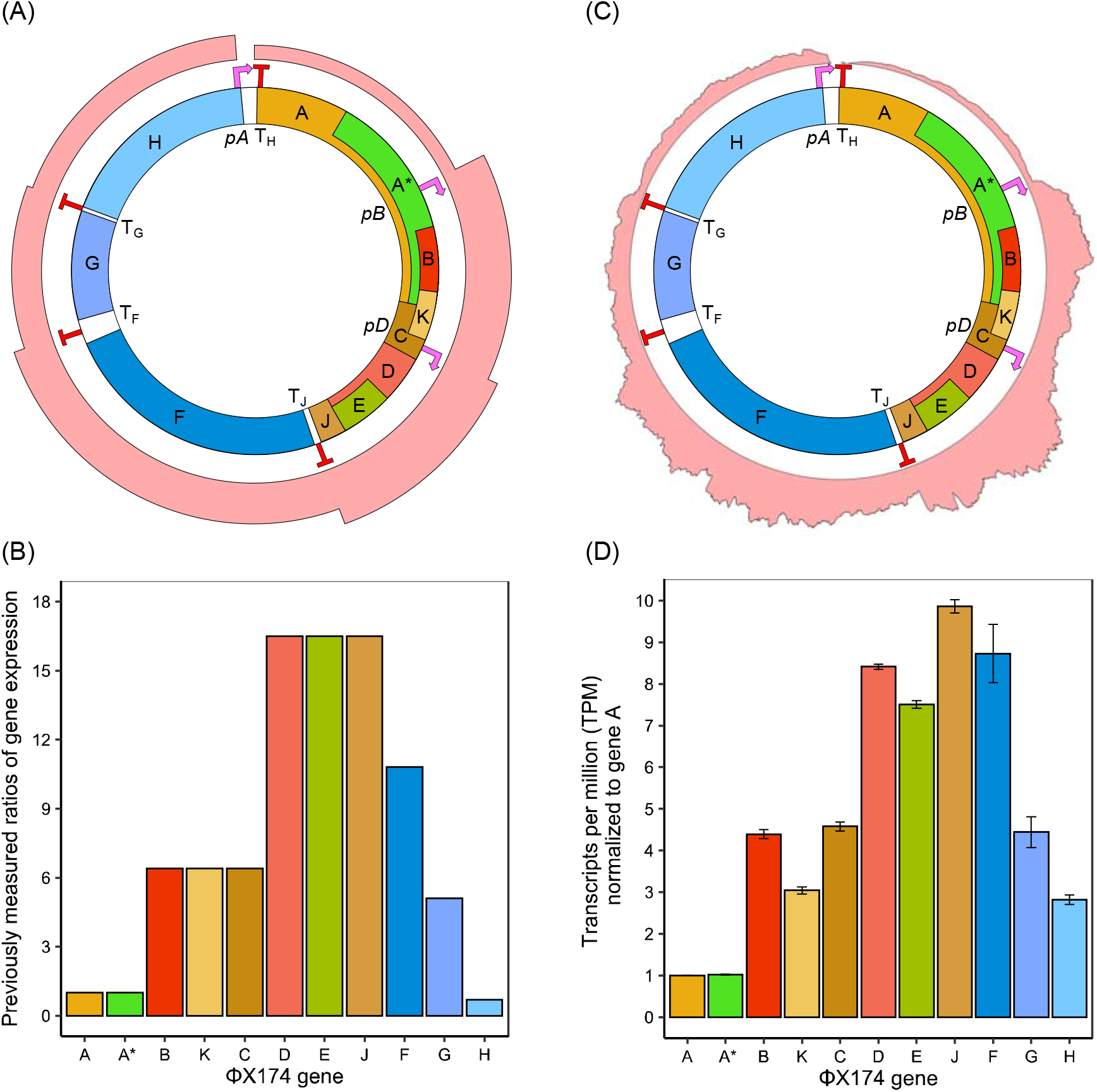
Models of ϕX174 transcription. (A) Previous model of øX174 transcription adapted from (Fane et al., 2005). Pink bars represent idealized mRNA quantities with initiation occurring at each known promoter and a step-wise decrease in transcriptional current occurring at each Rho-independent terminator. Gene colors represent broad protein function with red for scaffolding proteins, blue for capsid proteins, yellow for viral propagation proteins, and green for host interacting proteins. Promoters are shown in pink and terminators in red. (B) Previous model of relative transcription of øX174 genes generated from qPCR data (Brown et al., 2010; Zhao et al., 2012). (C) RNA-seq measurements of øX174 infection. Read mapping density across the øX174 genome shown by pink bar. Trimmed reads from biological replicates (n=3) mapped to reference øX174 genome (Genbank no. NC_001422.1) using Geneious Prime software. (D) RNA-seq measurements of relative transcription of øX174 genes. Values are the average of three biological replicates with error bars showing one standard deviation. Transcripts per million calculated from CDS feature annotations using Geneious Prime.

The ϕX174 single-stranded positive (+) sense genome that is initially transferred to the *E. coli* cytoplasm upon infection is thought to initiate transcription only after the complementary, or antisense (−) strand is synthesized, producing the double-stranded replicative form genome (Supplementary Fig. S1) (Sinsheimer et al., 1968). ϕX174 transcription is thought to exclusively use the antisense (−) genome strand as template, therefore the mRNA transcript sequence is identical to the viral (+) DNA strand (Supplementary Fig. S1) (Hayashi et al., 1963).

Unlike other coliphages, such as T7, ϕX174 does not encode an RNA polymerase (RNAP), and as a result, is totally dependent on the *E. coli* host’s transcription system. The ϕX174 RNAP promoter locations were previously identified by a combination of binding assays (Chen et al., 1973) and hybridization mapping (Axelrod, 1976a, b), revealing three major promoters upstream of genes A, B, and D (Fig. 1A and Supplementary Table S1). Additional *in vitro* studies mapped RNAP binding to specific genomic locations using electron microscopy, and determined multiple additional binding sites: the 3` end of gene G, and weak binding at two locations in the 5` end of gene F (Rassart and Spencer, 1978; Rassart et al., 1979). Follow up studies using a supercoiled pKO-1 plasmid and galactose activity to validate the identified promoter sites *in vivo* showed different relationships between *pA*, *pB*, and *pD* depending on the normalization methods employed (Ringuette and Spencer, 1994; Sorensen et al., 1998) (Supplementary Table S1). Furthermore, sequence context beyond the −35 and −10 regions seemed to have dramatic effects on promoter strengths, indicating that native ϕX174 transcription was affected by mRNA stability or enhancement sequences which were not transferred to their reporter plasmid (Sorensen et al., 1998). These context-specific effects are also consistent with prior findings (Hayashi and Hayashi, 1985; Hayashi et al., 1989) showing that better *in vivo* measurements of transcription from the øX174 genome itself are needed to address these inconsistencies.

The current understanding of øX174 transcriptional termination is that there are four Rho-independent (intrinsic) terminators located in the intergenic regions between genes J-F, F-G, G-H, and H-A, named T_J_, T_F_, T_G_, and T_H_ respectively (Fig. 1A) (Hayashi et al., 1981; Hayashi et al., 1989). However, past work shows a potentially more complex picture where *in vitro* experiments detected a single Rho-dependent terminator in gene A (Axelrod, 1976a, b) and multiple Rho-dependent terminators in gene F (Axelrod, 1976a, b; Kapitza et al., 1979).

Taken together, the contemporary ϕX174 transcriptome model is that it is driven by three constitutive promoters (*pA*, *pB*, and pD) of increasing strength and regulated by a set of four terminators (T_J_, T_F_, T_G_, and T_H_) with efficiencies below 100%, resulting in a series of transcripts of various lengths, tiling across the genome and forming six distinct regions with varying total transcript abundance (Fig. 1A). Recently, qPCR measurements of the relative transcript abundance in each of the six regions showed ratios of 1:6:17:11:5:1 for A:B:D:F:G:H transcripts, respectively (Fig. 1B) (Zhao et al., 2012). These results correspond well with existing knowledge of ϕX174 transcription, however the methods employed were only able to determine broad transcription trends within the phage and did not provide high resolution mapping of the transcriptome.

In this work we use next-generation RNA sequencing to map ϕX174 transcription for the first time at base-pair resolution. We address a number of open questions from the ϕX174 corpus and find evidence for a Rho-dependent terminator, and one new promoter candidate. Additionally, we identify several new missense single-nucleotide polymorphisms within structural genes. Surprisingly, we also measure small amounts of antisense RNA that may indicate previously undiscovered transcriptional activity in this phage.

## RESULTS

In order to create a base-pair resolution map of the øX174 transcriptome we introduced wild-type ϕX174 virions to *Escherichia coli* NCTC122 (C122) cultures at a multiplicity of infection (MOI) of 5. After 20 mins of growth, corresponding to the point just before lysis, still-intact cells were harvested and RNA isolated from them. The RNA was subsequently used to produce Illumina libraries and sequenced, followed by mapping to the ϕX174 genome. Read coverage was very deep with every base of the øX174 genome covered by between 900 – 318,502 separate reads. The resulting sequence reads showed a wide range of abundances across the genome that varied in magnitude in a manner that agreed well with previously annotated promoters and terminators (Fig. 1C).

We next compared gene-specific read abundance using the Transcript per Million (TPM) measurement which normalizes against both gene length as well as sequencing depth within each sample (Wagner et al., 2012). The results of this analysis (Fig. 1D) showed strong correspondence overall to previous measurements of øX174 gene expression (Fig. 1B) (Zhao et al., 2012). In contrast to the overall trend, the expression of genes K and E in our measurements were lower than expected by 53 % and 55 %, while genes J, F, and H were higher than expected by 40 %, 19 %, and 403 %, respectively (Fig. 1D).

### Quantifying promoter and terminator activity

To measure the relative strength of øX174 promoters *pA*, *pB*, and *pD* we compared TPM counts upstream and downstream of each known promoter site. The results of this analysis revealed that *pB* and *pD* increased transcriptional current, the flux of RNA polymerases passing a given point and generating RNA transcripts (Bonnet et al., 2013), by 2.0 and 2.5 fold, respectively (Fig. 2A). Measurement of *pA* activity was complicated by the fact that there is a promoter-terminator overlap within the pA-TH region, where the end of the promoter sequence is immediately followed by the beginning of the terminator stem (Supplementary Fig. S2). We observed a net loss of transcriptional current across the pA-T_H_ feature (Fig. 1C), which we interpret as a result of fewer RNAP initiation events from *pA* than termination events by T_H_. Therefore, our calculated strength for *pA* was below 1 (Fig. 2A).

**Fig. 2.**
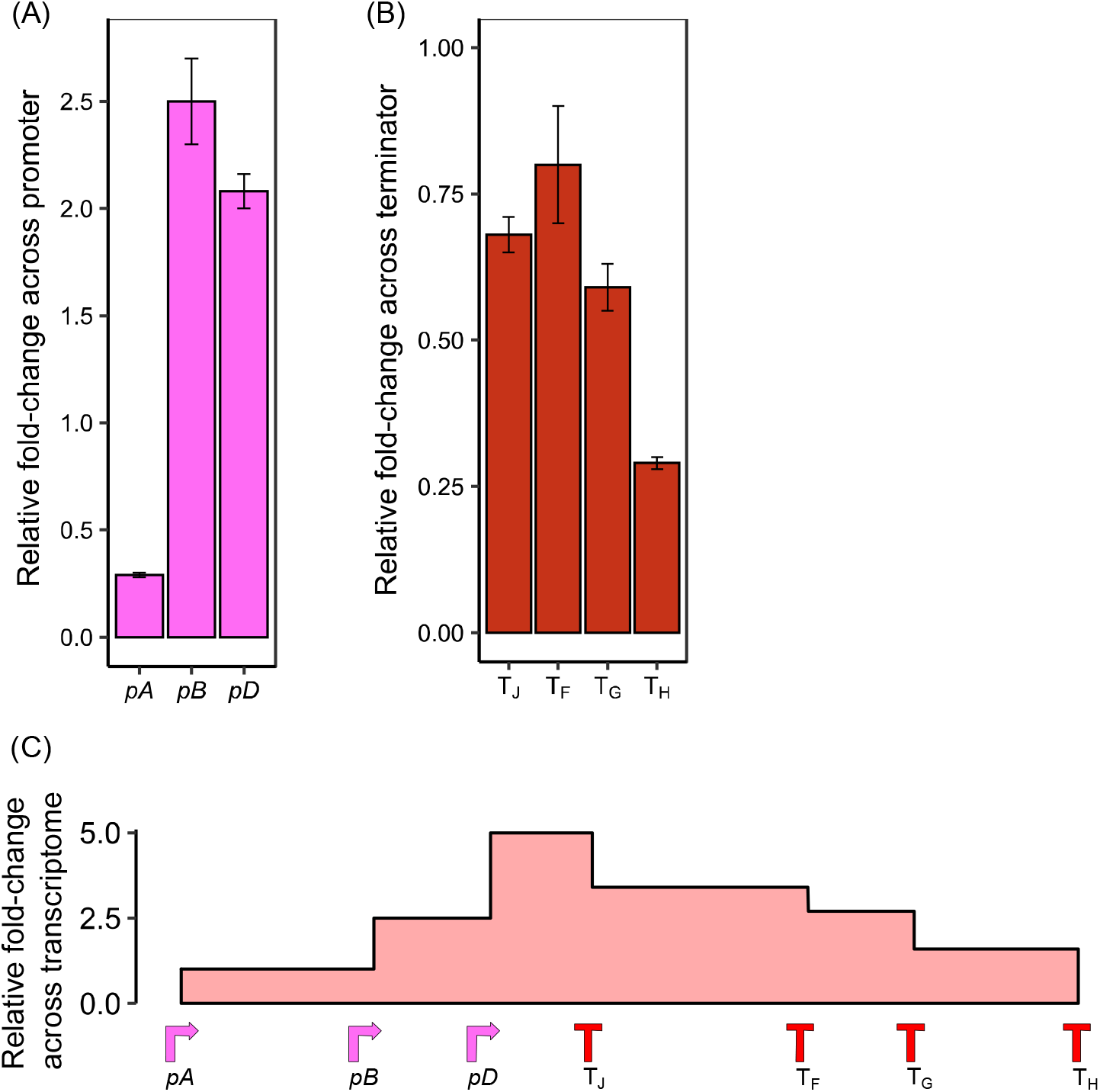
Measurements of canonical promoter and terminator strengths across øX174 genome using RNA-seq measurements. (A) Relative promoter strengths. (B) Relative terminator strengths. (C) Cumulative transcriptional current from combined promoter and terminator activities. Promoter and terminator strengths were measured by calculating differences in transcripts per million (TPM) read counts 100 bp downstream versus upstream of regulatory feature using Geneious Prime. Values are the average of three biological replicates with error bars showing one standard deviation.

Using the same analysis method, we measured the reduction in transcriptional current across the T_J_, T_F_, T_G_, and T_H_ terminators. This analysis showed that terminators T_J_, T_F_, and T_G_ only blocked 20-40% of transcriptional current flowing through them, whereas T_H_ was more effective, blocking 70% of transcriptional current (Fig. 2B). The *in silico* RNA folding energies of the terminator sequences do not correlate with measured termination efficiency (Supplementary Fig. S3).

Together, our measurements of promoter and terminator strengths update the quantitative description of the cumulative network that controls øX174 transcription (Fig. 2C).

### Determining presence of cryptic promoters and terminators

To determine whether there may be any undiscovered promoter elements within the øX174 genome, we identified areas where read abundances increased without a known promoter upstream. For example, the middle of gene E and the 5’-end of gene F (Fig. 1C). Next we used two computational tools to generate a series of predicted promoter sites and then analysed our data to determine if any of these promoters occurred in locations close to the transcription increases (Supplementary Fig. S4A). These tools were able to predict *pB* and *pD* but not *pA*, likely due to the overlap with T_H_ (Supplementary Fig. S2). One new promoter candidate (btss49) was identified at the 5` end of gene C (Fig. 3A) and preceded a noticeable increase in transcription (Fig. 3C), despite a modest measured strength due to local read coverage noise (Fig. 3B). The predicted btss49 promoter was identified as a σ24 type, which is generally involved with heat shock, and extracytoplasmic and envelope stress in bacteria (Rhodius et al., 2006). However, a detailed analysis of the potential transcription start site (TSS) revealed that while σ24 promoters typically have 5–6 nt between the –10 sequence and the +1 TSS (Rhodius et al., 2006), we see a 23 nt stretch between btss49 and the start of transcriptional increase (Fig. 3C). This discrepancy suggested to us that there was either degradation of the 5’ end of the btss49-initiating transcript or that while btss49 was predicted by bTSSfinder, another non-predicted promoter sequence may also or instead be in the same genomic region. Manual inspection of the sequence surrounding btss49 revealed a possible σ70 promoter sequence 40 nucleotides upstream of the TSS (as judged by RNA-seq read increase): AAGC<u>TCTTAC</u>TTTGCGACCTTTCGCCA<u>TCAACT</u>AACGATT, with potential −35 and −10 sequences conforming to the σ70 consensus underlined.

**Fig. 3.**
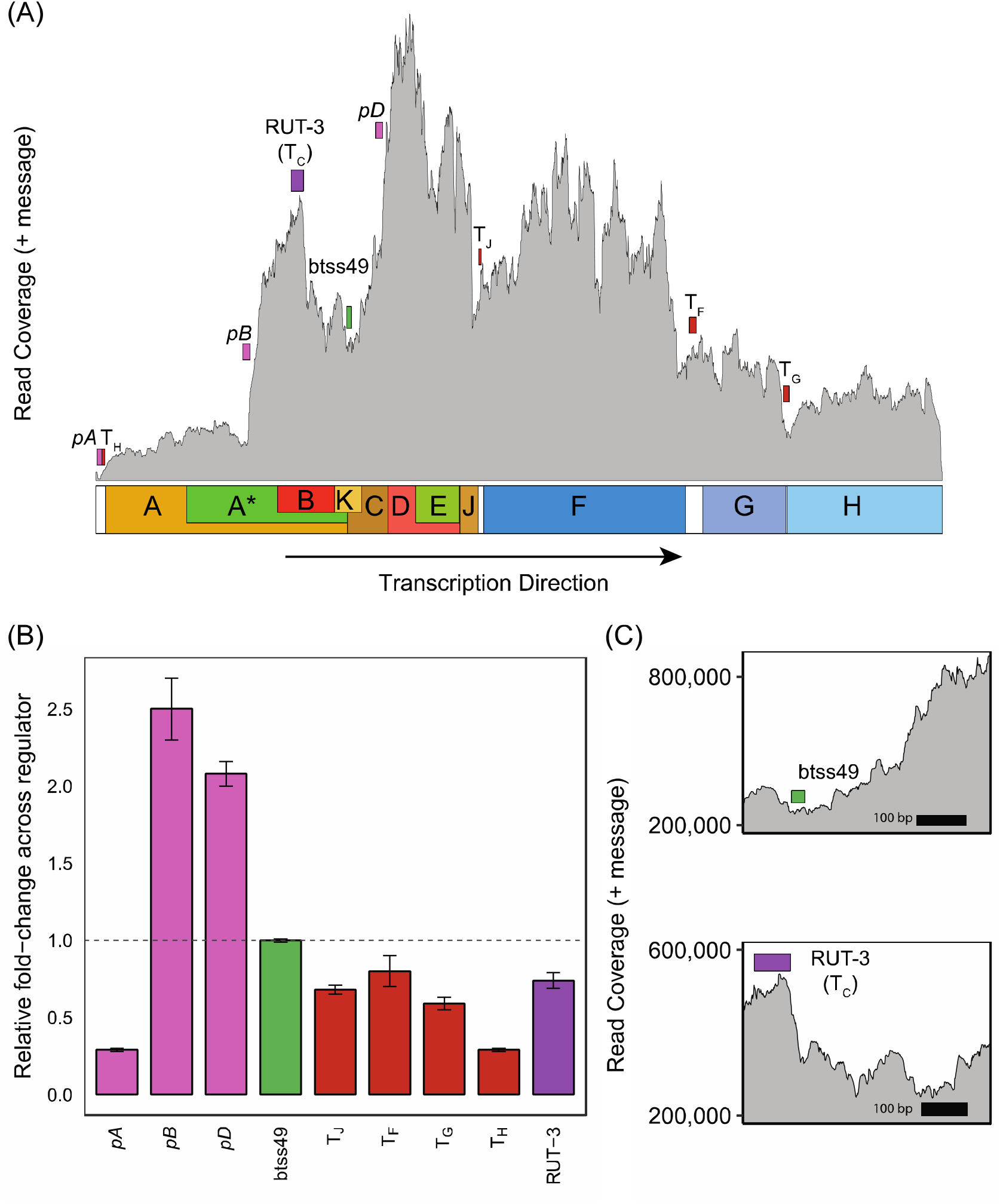
Computationally identified putative regulatory elements in øX174 genome. (A) Location of canonical and putative regulatory elements overlaid onto average read counts mapped across øX174 transcriptome (grey). Canonical promoters (pink), terminators (red), predicted promoter btss49 (green), and putative Rho-dependent terminator RUT-3 (T_C_) (purple) are annotated to contain total sequence length of feature. (B) Canonical and putative regulator strengths. Strengths were measured by calculating differences in transcripts per million (TPM) read counts 100 bp downstream versus upstream of the feature using Geneious Prime. Values are the average of three biological replicates with error bars showing one standard deviation. (C) Detailed view of predicted btss49 promoter and RUT-3 terminator.

We next looked for cryptic terminators by identifying regions where transcriptional current is reduce without the presence of one of the four canonical terminators. For example, the 5’-end of genes B and E (Fig. 3A). We used three computational tools to predict the presence of Rho-independent terminator locations within the øX174 genome. Despite these tools detecting all known terminators T_J_, T_F_, T_G_, and T_H_, they did not find any additional terminators mapping close to a sharp reduction in read abundance (Supplementary Fig. S4B), leading us to conclude that under our experimental conditions there are no Rho-independent terminators within øX174 left to be discovered.

We next looked for Rho-*dependent* terminators in the øX174 transcriptome. Rho-dependent terminators have been observed before *in vitro* (Axelrod, 1976a; Brendel, 1985; Smith and Sinsheimer, 1976a, b, c) but failed to be substantiated when probed *in vivo* (Hayashi et al., 1981). To identify locations of potential Rho utilisation (RUT) sites, which precede the Rho-terminator region by 10-100 nt (Koslover et al., 2012; Ray-Soni et al., 2016), we used the RhoTermPredict algorithm (Di Salvo et al., 2019). This analysis identified 13 RUT sites across the øX174 genome (Supplementary Fig. S5). Two of the predicted RUT sites, RUT–7 and RUT– 8, correspond with previously observed Rho-termination sites within gene F (Kapitza et al., 1979), however, no decrease in transcriptional current across the features was observed under the growth conditions tested (Supplementary Fig. S5).

A third site, RUT-3, corresponded broadly with the previously predicted location of T_C_, a putative Rho-dependent termination site which was previously reveal from *in vitro* assays (Smith and Sinsheimer, 1976a, b, c). We observed that the transcriptional current decreased significantly within 100 nt downstream of the predicted RUT-3 feature (Fig 3A and 3C). We measured the termination strength of RUT-3 and found it is comparable to other terminators within the øX174 genome with a 0.74 fold change occurring across the region containing the RUT site (Fig. 3B) and within 100 nt downstream (Fig. 3C). This analysis leads us to conclude that the RUT-3 Rho-dependent terminator is likely an active transcriptional terminator within øX174 under our laboratory conditions.

### Single Nucleotide Polymorphism

We next analysed the RNA-seq data to identify sites in the øX174 transcriptome containing single nucleotide polymorphisms (SNPs) within the measured phage population. Three SNPs were identified, all of which result in amino acid changes in their respective proteins (Table 2). The mutations occurred at different frequencies, with the gene F mutation (1347A>G) found at a frequency between 50.5-52.2 % of reads across the three biological replicates, while two gene H mutations (3112T>C, 3132G>A) were only found at 20.3-21.1 % and 40.2-41.9 % frequency, respectively.

**Table 2.**
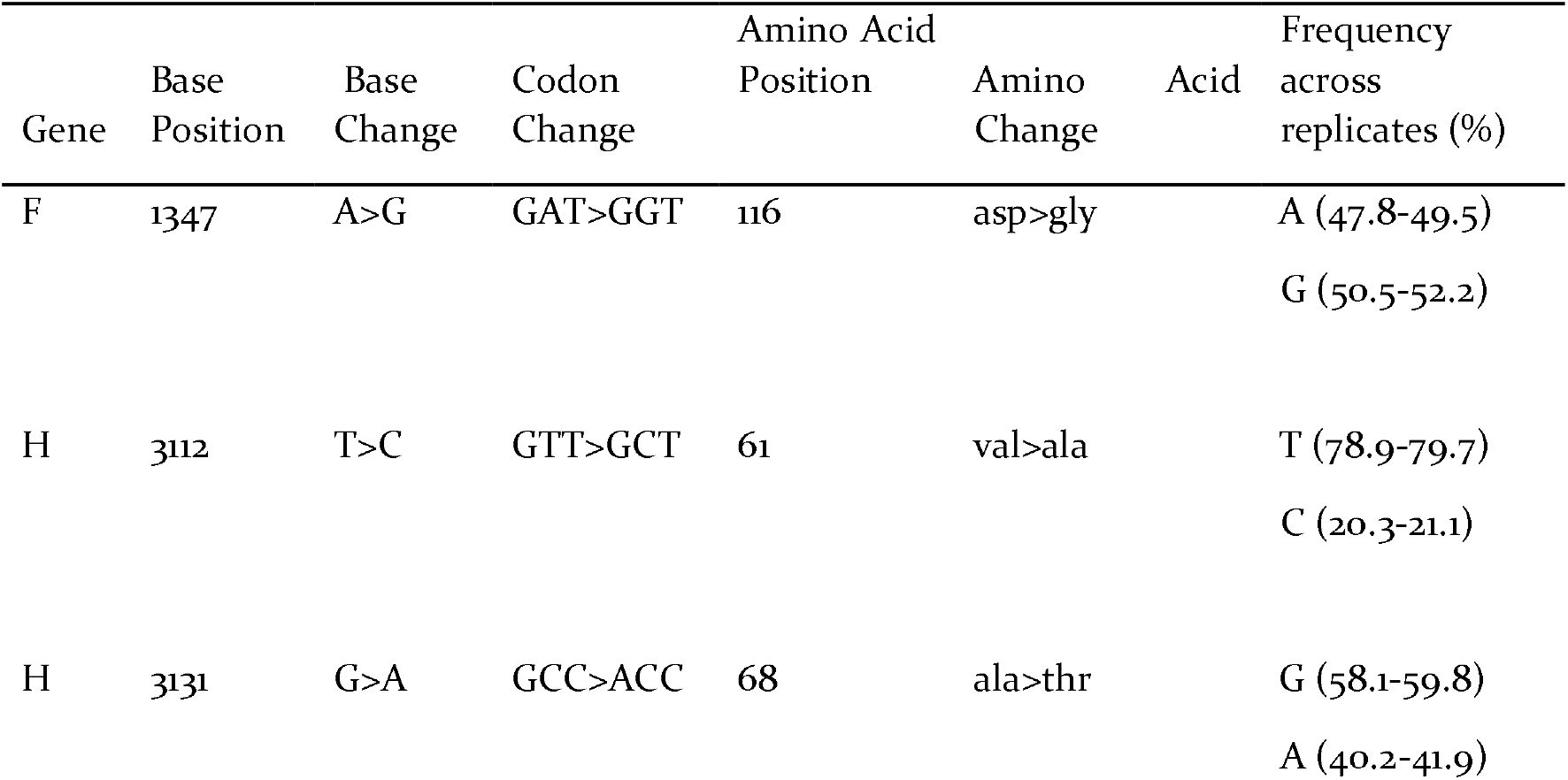
Single-nucleotide polymorphisms observed within øX174 population.

| Gene | Base Position | Base Change | Codon Change | Amino Acid Position | Amino Acid Change | Frequency across replicates (%) |
| --- | --- | --- | --- | --- | --- | --- |
| F | 1347 | A>G | GAT>GGT | 116 | asp>gly | A (47.8-49.5)<br>G (50.5-52.2) |
| H | 3112 | T>C | GTT>GCT | 61 | val>ala | T (78.9-79.7)<br>C (20.3-21.1) |
| H | 3131 | G>A | GCC>ACC | 68 | ala>thr | G (58.1-59.8)<br>A (40.2-41.9) |

### Discovery of antisense (−) RNA transcripts

Previous experiments seeking to identify all øX174 transcripts failed to detect the presence of any antisense RNA messages and predicted that if any messages were to be found in the future, they would constitute <5 % of total reads (Hayashi et al., 1963). The sequencing method we used in this work is designed to enable identification of which strand each sequencing read originates from. Surprisingly, during our analysis we mapped between 0.21%−0.38% of the total reads from each biological replicate (Supplementary Table S2) across the entire antisense (−) strand (Fig. 4A). The pattern of the antisense reads seemed to qualitatively match that of the sense strand reads, possibly due to strand mismarking during library preparation. Interestingly, several areas of large deviation from the sense strand read abundance pattern were also seen within genes G, H, and A (Fig. 4A), which indicated to us that we could also be measuring low abundance authentic antisense reads.

**Fig. 4.**
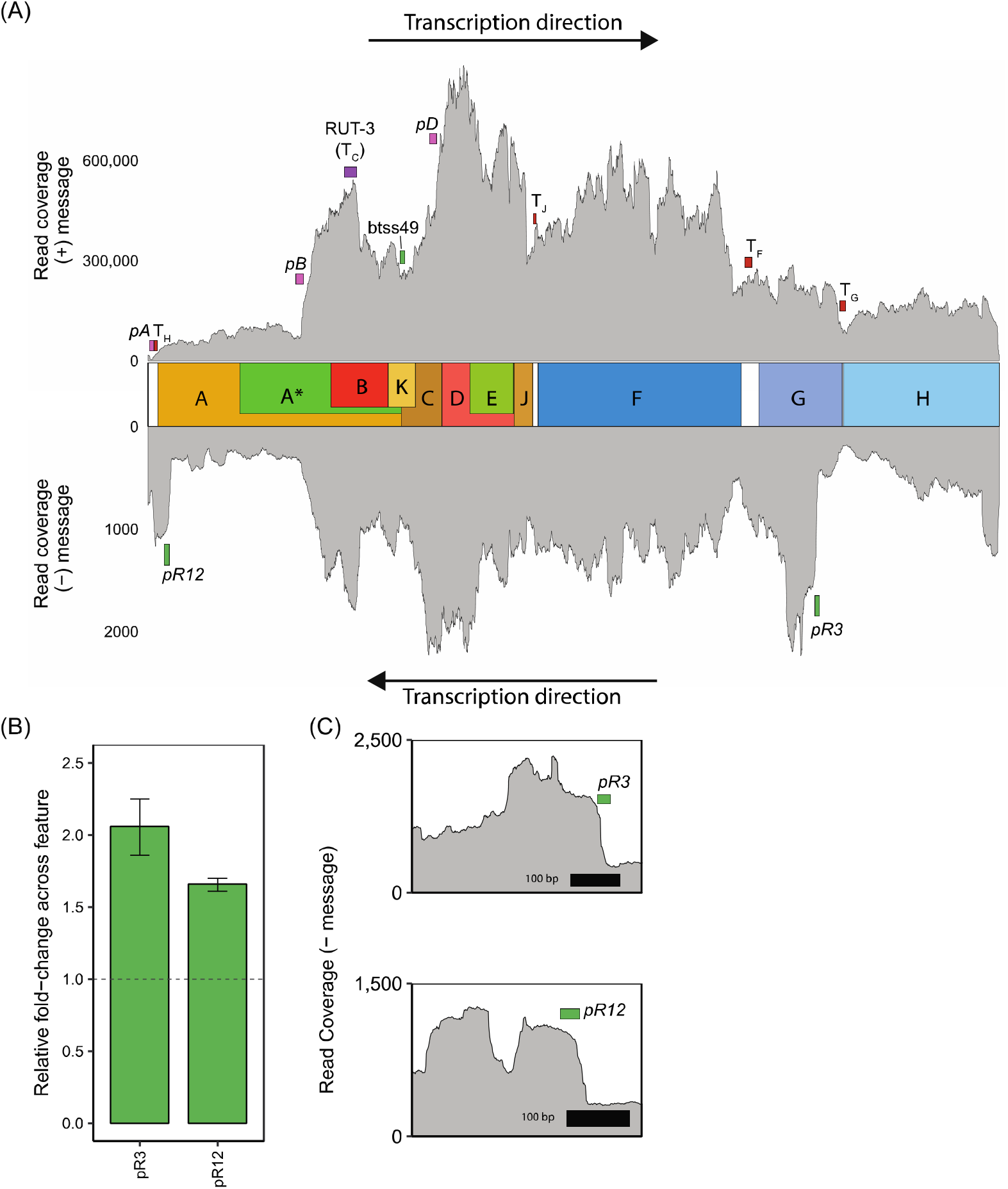
Comparison of sense (+) and antisense (−) read mapping with putative transcriptional control elements. (A) Sense (top) and antisense (bottom) reads aligned to the øX174 genome. Predicted antisense (−) promoters (green) are shown along with canonical and predicted sense (+) strand regulatory elements. Colored boxes represent the whole annotated sequence predicted from respective software. (B) Putative antisense promoter strengths measured by calculating differences in transcripts per million (TPM) read counts in 100 bp windows downstream versus upstream of each predicted promoter feature. (C) Detailed view of predicted pR3 and pR12 promoters.

To investigate this antisense RNA phenomena further, we used computational tools to identify any predicted promoter and terminator elements encoded on the øX174 sense (+) strand that could be responsible for driving the production of antisense RNA. We detected 12 potential promoters spread across the entire genome (Supplementary Fig. S6A and Supplementary File S2), but only pR3 and pR12 were located upstream of large increases in transcriptional current (Fig. 4C) that does not mirror the sense read pattern (Fig. 4A).

We also identified 11 potential terminator elements on the antisense (−) strand (Supplementary Fig. S6B), however none were adjacent to transcriptional current declines.

Next, we measured the strength of pR3 and pR12 putative promoters and found transcriptional current to increase across the feature at 2.06 and 1.66-fold, respectively (Fig. 4B). We conclude these newly discovered promoters may be responsible for the small number of antisense reads observed in our experiments that cannot be attributed to sense RNA mismarked as antisense during library preparation.

## DISCUSSION

Bacteriophage øX174 has been studied since the dawn of molecular biology, but our understanding of øX174 transcription has not kept pace with recent improvements in sequencing technology. In this work, we sought to update the øX174 transcriptome model by generating a new high-resolution map using next-generation sequencing technology. We used this high-resolution map to identify and quantify the strength of known regulatory elements, identify potentially new regulatory elements, as well as to identify novel antisense transcription and corresponding regulatory elements.

### Quantification of øX174 gene expression

From our analysis we were able to broadly confirm relative øX174 gene expression patterns but observed that RNA abundance decreased significantly within the regions immediately after initiation at *pB* and *pD* (Fig. 1C), which was not captured in previous transcription models (Fane et al., 2005; Zhao et al., 2012). Whereas previous models assumed that little to no variation occurred outside of known regulatory events, we were able to detect a distinct decrease in read coverage at the 3` end of gene B, which resulted in a decrease in measured gene K expression compared to literature (Fig 1B and Fig. 1D).

The variation we detected between our TruSeq RNA-seq data and previous TaqMan qPCR data (Zhao et al., 2012) is likely a result of the known differences in sensitivities between the two approaches (Schuierer et al., 2017). For example, our RNA-seq results to previous qPCR results, shows genes J, F, and H being more abundant by 40 %, 19 %, and 403 % respectively (Zhao et al., 2012). Differences in measured gene expression may also be due to differences in quantitation methods, or the MOI used in the respective experiments.

However, the 53% reduction in gene K expression we observe is not explained by either of these reasons and appears to be a result of the Rho-dependent terminator RUT-3 (T_C_) which we have identified and characterized in this work (Fig. 3C).

### Measurements of canonical promoters *pA*, *pB*, and *pD* strengths

We quantified the effect each øX174 promoter has on the transcriptional current and found that *pB* and *pD* increases transcriptional current by 2.5 and 2.0-fold, respectively. The combined pA-T_H_ element actually has higher termination activity than initiation activity, resulting in a net 0.29-fold transcriptional current change from the 3’-end of gene H to the 5’-end of gene A (Fig. 1C and Fig. 2A). The rank order of the promoter strengths agrees with previous work but qPCR measurements showed 6.4, 2.5, and 0.7-fold changes for *pB*, pD, and pA-T_H_, respectively (Zhao et al., 2012).

Compared to other bacterial and phage promoters, øX174 promoters are significantly weaker. For example, multiple common inducible promoters, *pBAD*, *pBAD2*, *pTAC* increase transcriptional current 8, 9, and 10-fold, respectively (Gorochowski et al., 2017), whereas the ϕX174 promoters are in the range of 2.0 - 2.5-fold (Fig. 2A). When cataloguing constitutive promoters in the bacterium *Bifidobacterium breve*, which uses σ70 promoter sequences similar to *E. coli*, promoter strengths were calculated using RNA signal/gDNA baseline. This determined relative transcriptional strengths significantly above that of the common inducible and ϕX174 promoters with fold changes of 150 – 21,000, with the vast majority between 150 and 1,100 (Bottacini et al., 2017). Similarly, an analysis of a series of promoters in *Zymomonas* mobilsis found a promoter fold change range of 24 – 25,000 (Yang et al., 2019). A similar approach was taken with the T7 promoter, where the strength was calculated at 169 fold (Komura et al., 2018).

No standard characterisation methods exists for using RNA-seq to characterize promoters, creating difficulties in making a direct comparison to ϕX174 promoters. The majority of prokaryotic promoters transcribe isolated genes allowing for a comparison of RNA to DNA counts, whilst our system has overlapping transcriptional units requiring us to calculate relative changes in RNA counts. However, the low strength of the ϕX174 promoters is likely explained by high number of phage particles per cell, reducing the requirement for high transcription activity.

### Measurements of known terminator T_J_, T_F_, T_G_, T_H_ strengths

øX174 transcription is known to generate a range of mRNA species due to terminator read-through (Hayashi et al., 1981). Using our high-resolution sequencing data we measured for the first time, to our knowledge, the termination efficiency of each canonical terminator by comparing read depth before and after each terminator. Our measurements show that T_J_, T_F_, and T_G_ are all relatively poor Rho-independent terminators, reducing the transcriptional current that passes through them by between 0.80 and 0.59-fold (Fig. 2B). These values put these three øX174 terminators in the lowest strength quartile of all *E. coli* terminators, on par with the *endA* and *araBAD* terminators (Chen et al., 2013; Greinert et al., 2012; Jekel and Wackernagel, 1995; Lee et al., 1986). Although measurements of the strength of the T_H_ terminator were confounded by the overlapping *pA* promoter, the combined element is more than twice as effective at terminating transcriptional current than the next strongest terminator T_G_, with the calculated terminator strength placing it in the second lowest strength quartile of *E. coli* promoters (Chen et al., 2013; Greinert et al., 2012; Jekel and Wackernagel, 1995; Lee et al., 1986).

Although relative weak individually, taken together as an entire system, the øX174 T_J_, T_F_, and T_G_ terminators in series have a large combined effect on transcriptional current with only ~ 18 % of transcriptional current remaining by the time polymerases reach pA-T_H_ (Fig. 2C). The pA-T_H_ element effectively ‘resets’ transcriptional current by further reducing the current to 9 % of its peak in time for the start of gene A.

### Detecting novel promoters

From our RNA-seq dataset in this work we observed several areas in the transcriptome where read depth rose in the absence of any canonical promoter. Using computational tools we identified one new candidate promoter preceding *pD* by 157 base-pairs in a region where transcription current increases by over 3-fold over 350 base-pairs (Fig. 3C).

Interestingly, the novel promoter btss49 was identified as a σ^24^ type which works in concert with the σ^32^ sigma factor and together are activated under extreme cellular heat shock and capsule and membrane disruption (Raina et al., 1995). Previously, a possible σ^32^ promoter was identified close to the canonical *pB* promoter in experiments subjecting øX174 to heat shock during infection (Zhao et al., 2012). Both σ^24^ and σ^32^ are known to be activated during bacteriophage infection and associated stress response (Bahl et al., 1987; Osterhout et al., 2007). Together, it seems highly likely that øX174 uses some additional promoters that are activated by the stress caused by a phage infection, and corresponding membrane disruption, to increase transcription of certain areas of the genome and increase fitness.

### Detecting novel terminators

To identify novel termination events in øX174 transcription, we performed computational analysis of both Rho-independent and Rho-dependent terminators and correlated these to falls in transcriptional current unexplained by canonical terminators. Previous work in ϕX174 suggested there could be additional terminators of both types still to be discovered (Axelrod, 1976a; Fane et al., 2006; Hayashi et al., 1988).

Interrogating our RNA-seq data we identified one Rho-dependent terminator (RUT-3) (Fig. 3A), co-located with a previously detected Rho-dependent terminator from past øX174 literature, T_C_ (Smith and Sinsheimer, 1976a, b, c). Our detection of RUT-3/T_c_ suggests that the earlier *in vivo* RNA characterisation methods were not sensitive enough to detect the relatively low level of activity at this regulator. We now name this Rho-dependent terminator T_C_ after the original name it was given (Smith and Sinsheimer, 1976a, b, c) and in keeping with the naming conventions currently used for øX174 terminators.

Rho-dependent termination is a factor-driven terminator method requiring the binding of a hexameric Rho protein complex to DNA, which translocates to the attached RNAP, forcibly ending transcription. Rho-dependent terminators are known to exist in bacteriophage P2, T5, and lambda (Brunel and Pilaete, 1985; Court et al., 1980; Linderoth and Calendar, 1991). Rho-dependent terminators in lambda phage are used to transition from early to late genes during its lifecycle (Casjens and Hendrix, 2015) and the lambda N protein suppresses both Rho-independent and Rho-dependent terminators (Nudler and Gottesman, 2002). Bacteriophage P4 is also known to produce the Psu protein which is an antiterminator of Rho-dependent terminators in both the P2 phage and *E. coli* (Linderoth and Calendar, 1991; Linderoth et al., 1997; Pani et al., 2006). With the presence of the T_C_ Rho-dependent terminator better established in øX174 it may be worthwhile to assess whether any existing øX174 gene product has antitermination activity.

### Detecting novel single nucleotide polymorphisms

Despite a large catalogue of genome sequence variants already identified within ϕX174 (Bull et al., 2003; Bull et al., 2000; Bull et al., 1997; Crill et al., 2000; Sackman et al., 2017; Wichman et al., 2005; Wichman et al., 2000), the three SNPs detected here in genes F and H have never been observed before in wild-type øX174 populations (Table 2), although the gene F asp>gly mutant was represented within a synthetic library created recently (Faber et al., 2020).

Previous studies investigating the effect of SNPs on ϕX174 fitness found an excess of mutations within gene A, and a deficit within the capsid proteins, genes F, G and H, even after correcting for differing gene lengths (Wichman et al., 2005). The missense mutations in genes F, G, and H accumulated in a near-linear fashion across the entire 13,000 generation experiment, however, it is not known if these capsid mutations were adaptive or neutral as no clear phenotypic advantage was noted.

The wild-type øX174 used in this study was created using synthetic DNA to match the original Sanger sequence (Jaschke et al., 2019) (Genbank No. NC_001422.1) and differs from commercially available preparations in at least nine bases (Jaschke et al., 2012; Smith et al., 2003). It is perhaps unsurprising that novel polymorphisms are observed in this study since øX174 has a mutation rate of 1.0 × 10^−6^ substitutions per base per round of copying (Cuevas et al., 2009) and the phage genome was created synthetically to match a catalogued sequence rather than as a product of a naturally selected population.

### Antisense (−) transcription in ϕX174

In this study we observed for the first time antisense (−) RNA within ϕX174 (Fig. 4 and Supplementary Table S2). Antisense RNA read coverage varied by 25-fold across the genome (Fig. 4A). This variance in abundance indicates that the source of the reads was unlikely to be øX174 replicative form DNA contamination during the sequencing library prep, which would be expected to produce relatively uniform read coverage across the contaminating nucleic acid sequence.

Another possible source of contamination could have been the PhiX Control v3 RFI DNA used within Illumina sequencers, which would also have been expected to produce uniform coverage. We specifically did not use this control during our sequencing runs to prevent cross-contamination, but it is possible that small amounts of this control DNA could have been present within the sequencer or reagents. We disproved this idea by comparing the consensus sequence of the mapped antisense (−) sequences to the wild-type øX174 sequence (Genbank No. NC_001422.1) and the PhiX Control v3 sequence, which differ from each other at five positions (587G>A, 833G>A, 2731A>G, 2793C>T, 2811C>T). In all cases our antisense (−) consensus sequence matched NC_001422.1 and not PhiX Control v3, indicating the reads were from our øX174 libraries and not Illumina control contamination.

During replication of the complementary (−) strand of the øX174 genome, the primosome synthesizes short 9-14 nt RNA primers using the viral (+) strand as template (Supplementary Figure S1). These RNA primers would have the same sequences as the detected antisense RNA in this work. We can discount this possibility though because such short sequences would be removed by during the read trimming process. Manual inspection of the mapped reads also shows no sequences below 30 nt in length.

A further possible explanation could be the strand marking protocol allows a small proportion of strand misidentification and that reads identified as antisense are not. In this case we would expect the antisense reads to perfectly match the pattern of the sense reads, which is the case for the majority of the antisense reads mapping across the øX174 genome in our experiment, with the exception of two areas around genes G, H, and A (Fig. 4A).

Within the G to A region, putative promoters pR3 and pR12 can be detected upstream of antisense read abundance increases (Fig. 4C). These σ^70^ promoters (Supplementary File S2) appear to be relatively strong as we measure them to increase transcriptional current by 2.0 and 1.7 fold, respectively. This places pR3 at a similar strength to *pB*. When considered in the context of broader *E. coli* promoters they are relatively weak when compared to commonly used inducible promoters (Gorochowski et al., 2017).

Despite our observation of antisense (−) transcripts, the fraction of total reads were less than 1%. Additionally, no past work has ever shown evidence for proteins produced from the antisense (−) transcript (Pollock et al., 1978). Recently, we tried to identify any additional open reading frames ≥ 60 bp initiating with *E. coli’s* most common start codons (Hecht et al., 2017) from both strands of øX174 using mass spectrometry (Jaschke et al., 2019). We could only detect one additional protein produced from the sense (+) mRNA and no proteins from the antisense (−) transcript. While we cannot rule out the possibility of protein produced from the antisense (−) transcript, further investigation using more sensitive mass spectrometry methods such as Parallel Reaction Monitoring (Peterson et al., 2012; Vincent et al., 2019) would be needed to establish the absence of translation from the RNAs detected in the current study. There may be uses, from an evolutionary perspective, for non-translated mRNA production (Yona et al., 2018). For example, many genomes have a spectrum of genic and non-genic transcribed sequences allowing for the development of protein-producing open reading frames over evolutionary timescales from either canonical or non-canonical start codons (Carvunis et al., 2012; Hecht et al., 2017).

### Implications of updated transcriptome model of ϕX174

In this work we used a high-resolution RNA-seq dataset to update the transcription model of bacteriophage øX174 in several important ways. We were able to broadly confirm the known model for ϕX174 transcription, but added refined transcriptional ratios between genes and revealed areas of previously uncharacterized transcriptional initiation and termination.

The precise step-wise activity of the promoters and terminators of øX174 are important for the phage lifecycle to create different levels of transcripts encoding each gene since the only known control mechanisms are genome concentration, transcription rate, and translation efficiency (Fane et al., 2005; Hayashi et al., 1988). Our RNA-seq results match well with the previously measured protein abundances with ratios of B:D:J:F:G:H being 60:240:60:60:60:12 (Fane et al., 2006).

In contrast though, we see that gene E RNA abundance is lower than expected. Despite falling within a region of overall high RNA expression, there appears to be some mechanism to reduce transcript levels for gene E (Fig. 3A). This effect may be related to the low concentrations of E protein required for host lysis, aided by a very weak ribosome binding site (Bläsi et al., 1990; Maratea et al., 1985).

The Rho-dependent terminator, T_C_, seems to function to reduce gene K transcripts (Fig. 3A). We speculate that the role of this added regulation of gene K, which modulates phage burst size (Gillam et al., 1985), is likely related to limiting burst size in a way that increases øX174 fitness.

Subsequent recovery of the transcriptional current after T_C_ by *pD* is augmented by the putative heat shock promoter btss49 (Fig. 3A and 3C). The novel promoter, btss49 or a nearby sequence, appears to be important in recovering gene C transcript abundance. Gene C is important in switching from stage II to stage III DNA synthesis which results in increased ssDNA packing into phage capsids using the host machinery and viral proteins A and J (Hayashi et al., 1988; Mukai et al., 1979). It is likely that this phage lifecycle switch exploits the host stress response from the infection to drive the formation of phage particles prior to lysis.

This work also shows evidence for the first time of antisense transcription in ϕX174, which has not been detected in any other member of the *Microviridae*. In fact, the closest related phage that is known to produce antisense RNA are the *Podophage* such as P22 (Casjens and Grose, 2016; Liao et al., 1987).

Antisense transcripts can either encode open reading frames that are translated into proteins, or they can play a role interfering with sense RNA translation by ribosome occlusion or RNAse targeting (Wade and Grainger, 2014). We have no prior evidence of translation from antisense RNAs in øX174 (Jaschke et al., 2019; Pollock et al., 1978), so it is unlikely that the observed antisense RNA encodes ORFs producing protein. Regulatory antisense RNAs have been observed in lambda phage (Krinke et al., 1991), bacteriophage 186 (Dodd and Egan, 2002), P22 (Liao et al., 1987), and T4 (Belin et al., 1987). Antisense RNA has been detected in phage ϕ29 (Mojardín and Salas, 2016) and AR9 (Lavysh et al., 2017) but no definitive function has been yet assigned.

Although no other ssDNA phages are known to encode antisense RNA, it is relatively common among ssDNA viruses of eukaryotes. For example, Bean golden yellow mosaic virus, Beet curly top virus, Maize streak virus, Tomato pseudo-curly top virus, Porcine circovirus 1, and Bacilladnavirus all produce antisense RNA that contains genes (Briddon et al., 1996; Gilbertson et al., 1991; Lazarowitz, 1988; Niagro et al., 1998; Stanley et al., 1986; Tisza et al., 2020).

A different role for the antisense transcription seen in this work could be attributed to controlling the production of proteins A and A*, which are known to be critically important for ϕX174 replication and packaging (Roznowski et al., 2020), but also seem to be only needed at very low quantities for successful infections (Jeng et al., 1970). In fact, overexpression of these genes has been seen to be deleterious towards the host, therefore controlling their expression may be crucial to ensuring early stages of DNA replication are completed successfully before lysis gene E is expressed (Colasanti and Denhardt, 1985). In our work we see several ways that this tightly controlled production of proteins A and A* occurs. In phage known to produce cis antisense RNA from transcribing the opposite strand to a gene, one mechanism of gene expression disruption is through transcriptional interference (TI) caused by the head-on collision of the RNAP on each strand (Callen et al., 2004; Shearwin et al., 2005). This could be one possible role for the pR12 promoter located near to *pA* on the opposite strand, where it could play a role in decreasing *pA* transcription initiation, or dislodging any RNAP passing through the TH site that initiated at *pB* or *pD* (Fig. 4A). Therefore, we speculate that *pA* activity is tightly controlled through several independent mechanisms to supress the expression of A and A* genes during early infection to ensure host survival until gene E expression and controlled lysis occurs.

In summary, we have updated and expanded the current transcriptome model for ϕX174 by confirming and characterizing the effect of previously known regulatory elements within the phage. Specifically, we have confirmed the presence of a previously ignored Rho-dependent terminator, T_C_, and discovered a potentially novel promoter, btss49. Finally, we have revealed the first evidence for antisense transcription in øX174, and more broadly, in any ssDNA phage.

## MATERIALS AND METHODS

### Bacterial strains and øX174 propagation

Overnight cultures of *Escherichia coli* NCTC122 (National Collection of Type Cultures, Public Health England) were grown in 15 mL centrifuge tubes (Westlab, #153-560) in an Infors MT Multitron pro shaker at 250 RCF rotating orbitally at 25 mm diameter at 37 ^o^C containing Lysogeny Broth (LB) Miller media with 2 mM CaCl_2_ (phage LB) (Jaschke et al., 2019; Rokyta et al., 2009). Overnight cultures were diluted with fresh media so all had an OD_600nm_ of 1 and then split 1:50 into a 500 mL flat-bottomed Erlenmeyer flask (Duran, #2121644). Cells were grown in triplicate at 250 RCF and 37 ^o^C until a mid-log growth of 0.7 OD_600nm_ was achieved. Cells were centrifuged at 8,000 RCF and 4 ^o^C, washed with 5 mL ice-cold HFB-1 starvation buffer (Fane and Hayashi, 1991) (60 mM NH_4_CL, 90 mM NaCl, 100 mM KCl, 1 mM MgSO_4_•7H_2_O, 1 mM CaCl_2_, 100 mM Tris Base, pH 7.4) twice with centrifugation. Cells were finally resuspended with 5 mL ice-cold HFB-2 buffer (60 mM NH_4_Cl, 90 mM NaCl, 100 mM KCl, 1 mM MgSO_4_•7H_2_O, 10 mM MgCl_2_•6H_2_O, 100 mM Tris base, pH 7.4), and split equally (2.5 mL) into two 15 mL centrifuge tubes on ice. Infected samples of NCTC122 were treated with a ϕX174 MOI of 5. Infected samples were incubated at 14 ^o^C for 30 min to allow phage attachment without genomic insertion. These were then supplemented with 22.5 mL of 37 ^o^C pre-warmed phage LB media in 250 mL flat bottomed Erlenmeyer flasks (Simax, #1632417106250) and grown at 250 RCF and 37 ^o^C. Bacterial growth parameters have been reported to the best of our knowledge conforming with the MIEO v0.1.0 standard (Hecht et al., 2018).

A 5 mL sample was taken from the triplicates after 20 min and transferred to ice-cold 15 mL falcon tubes and stored on ice. These were then centrifuged at 3,500 RCF for 5 min at 4 ^o^C, (Eppendorf, 5430 R), and resuspended in 200 μL of ice-cold 1x PBS and briefly vortexed to mix the cells. The samples then had a 2:1 ratio of RNAprotect (Qiagen: #76506) added to them and centrifuged at 5,000 RCF for 10 min, before the supernatant was removed and the samples stored at −80 ^o^C.

### in silico genetic element prediction

Promoter prediction was performed via web analysis tools BPROM (Salamov and Solovyevand, 2011) and bTSSfinder (Shahmuradov et al., 2017) using their default settings and full length sequences for ϕX174 (Genbank No. NC_001422.1).

Previous Rho-independent terminator sequences from literature (Brendel, 1985; Godson et al., 1978; Hayashi et al., 1981; Otsuka and Kunisawa, 1982) were analysed with RNA folding simulations using the NUPACK web-server with default settings (Zadeh et al., 2011). Lowest-energy structures were reported.

Potential Rho-independent terminators were identified using the following software packages: FindTerm (Salamov and Solovyevand, 2011), iTerm-PseKNC (Feng et al., 2018), and RibEx: Riboswitch Explorer (Abreu-Goodger and Merino, 2005). Data was analysed via 300 nt windows with 50 nt overlaps between windows. The iTerm-PseKNC and RibEx packages were used according to default settings, while FindTerm was modified to set the Energy threshold value to –12.

Potential Rho-dependent terminators were identified by analysing the whole ϕX174 genome (Genbank No. NC_001422.1) with the RhoTermPredict python script (Di Salvo et al., 2019). All predicted sequences are collected in Supplementary File S2.

### Next-generation sequencing sample preparation

RNA was purified from 5 mL *E. coli* culture following infection using the RNeasy Mini kit (Qiagen: #74106) according to the manufacturer’s instructions, with the optional DNase on-column digestion step (Qiagen: #79254). RNA concentrations were measured with the QUBIT RNA assay (Life Technologies: #Q32852). Sequencing library preparation was performed by Macrogen Inc (S. Korea). The rRNA from samples was depleted with a Ribo-Zero Kit (Illumina) and the RNA library generated with a TruSeq Stranded mRNA kit for microbes. Sequencing was carried out with an Illumina HiSeq 2500 instrument in 2×100bp mode. PhiX Control Library was not used during the sequencing run to avoid cross-contamination.

### Sequence analysis

Raw reads were trimmed using Geneious Prime’s (Version 2019-1-3) implementation of the BBDuk trimmer (Bushnell, 2019) using the following command: java -ea -Xmx6000m -cp…/current jgi.BBDuk2 ktrimright=t k=27 hdist=1 edist=0 ref=adapters.fa qtrim=rl trimq=30 minlength=30 qin=33 in=input1.fastq out=output1.fastq. Following trimming, sets of paired reads from three biological replicates were mapped to the øX174 reference sequence (GenBank No. NC_001422.1) using the Geneious assembler (Version 2019-1-3) with the following settings: medium sensitivity, iterate: up to 5 times, minimum mapping quality: 30, and map multiple best matches: to none.

Following assembly, coding sequence (CDS) feature annotation expression levels were calculated using the ‘Calculate Expression Levels’ function of Geneious Prime with default settings with Ambiguously Mapped Reads: Count as Partial Matches.

Change in transcriptional current across promoter and terminator sequences were calculated by creating 100 bp CDS feature annotations up and downstream of the promoter/terminator followed by applying the ‘Calculate Expression Levels’ function (Geneious Prime) to those features using the Ambiguously Mapped Reads: Count as Multiple Full Matches setting.

SNPs were analysed through the ‘Find Variations/SNPs’ function of Geneious Prime with default settings.

Antisense (−) RNA was discovered by mapping trimmed ‘Read 1’ and ‘Read 2’ reads to the ϕX174 reference sequence. When Read 1 maps in the Forward direction or Read 2 maps in the Reverse direction, they are expected to be the identical sequence to the antisense (−) transcript (Illumina TruSeq Stranded mRNA Reference Guide (1000000040498 v00)). Between 0.21% - 0.38% of Reads 1 and 2 mapped in this way across the three biological replicates (Supplementary Table S2). Further analysis, such as promoter and terminator strength measurements were performed on this subset of the reads.

## ACKNOWLEDGEMENTS

We recognize that this research was conducted on the traditional lands of the Wattamattagal clan of the Darug nation. We thank Varsha Naidu, Liam Elbourne, and Sasha Tetu for helpful discussions and feedback.

## FUNDING

PRJ was supported by the Molecular Sciences Department, Faculty of Science & Engineering, and the Deputy Vice-Chancellor (Research) of Macquarie University. DYL is a recipient of the Macquarie University Research Excellence PhD scholarship (MQRES) and CSIRO PhD Top-up Scholarship (Program in Synthetic Biology).

## CONFLICTS OF INTEREST

The authors declare no competing conflicts of interest.

## APPENDIX A. SUPPLEMENTARY MATERIAL

Supplementary Material File S1: Fig. S1 – S6 and Tables S1-S2, PDF

Supplementary File S2: Computationally Predicted Promoter and Terminator Sequences.xlsx

