## Supplemental Material File S1 for "A high-resolution map of bacteriophage øX174 transcription"

**CONTENTS**

SUPPLEMENTAL FIGURES

Figure S1 DNA replication and transcription from sense and antisense strands of øX174

Figure S2 Structure of *pA*-T_H_ region

Figure S3 Predicted structures of transcriptional terminators within øX174 genome

Figure S4 Computationally predicted φX174 genetic elements regulating sense transcription

Figure S5 Computationally predicted Rho-Utilisation (RUT) sites

Figure S6 Computationally predicted φX174 genetic elements regulating antisense transcription

SUPPLEMENTAL TABLES

Table S1 Previously predicted sequences of φX174 promoters

Table S2 Stranded read mapping against NC_001422.1 reference sequence

SUPPLEMENTAL REFERENCES

SUPPLEMENTAL FILES

File S2 Promoter and terminator sequences

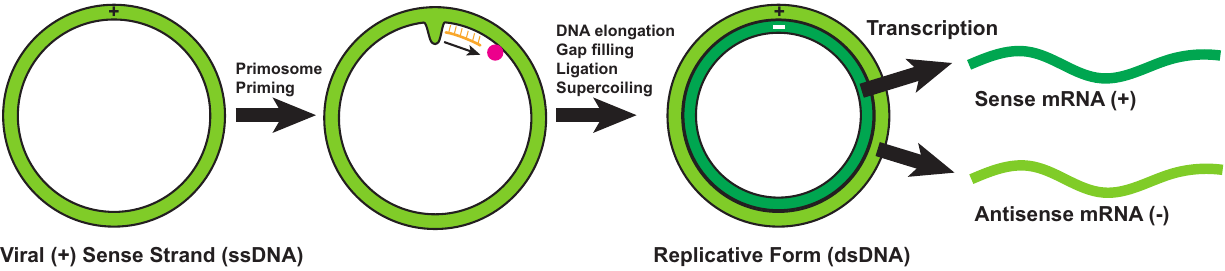

**Figure S1** DNA replication and transcription from sense and antisense strands of øX174. The viral (+) sense strand bulge at the gene F-G intergenic region forms a site for primosome assembly. The primosome synthesizes short 9-14 nt RNA primers along the length of the øX174 genome using the viral (+) sense strand as template. RNA primers prime the synthesis of the complementary (-) strand followed by gap filling, ligation, and supercoiling to form the replicative double-stranded DNA form the øX174 genome. Transcription using the complementary (-) strand of the genome as template results in the canonical sense mRNA (+) carrying all the known øX174 gene coding sequences. Transcription from the sense (+) viral strand of øX174 genome would result in antisense mRNA (-) with sequence identical to the complementary (-) strand. The RNA primers synthesized from the primosome would have the same sequence as antisense mRNA (-). (Hayashi et al., 1988)

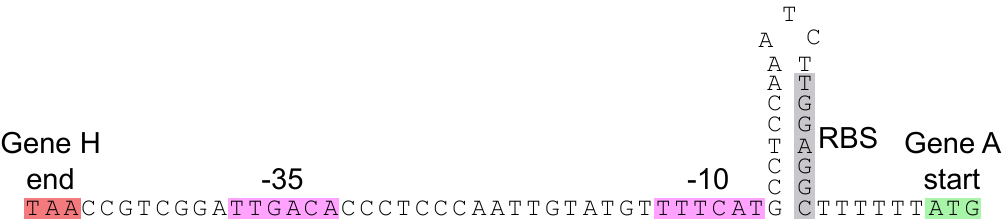

**Figure S2**. Structure of *pA*-T_H_ region. The H-A intergenic region contains multiple overlapping transcriptional regulatory elements as the promoter elements for *pA* are upstream to the stem-and-loop of T_H_. The stem-and-loop also contains the transcription start site as well as the ribosome binding site for gene A. Figure adapted from (Godson et al., 1978).

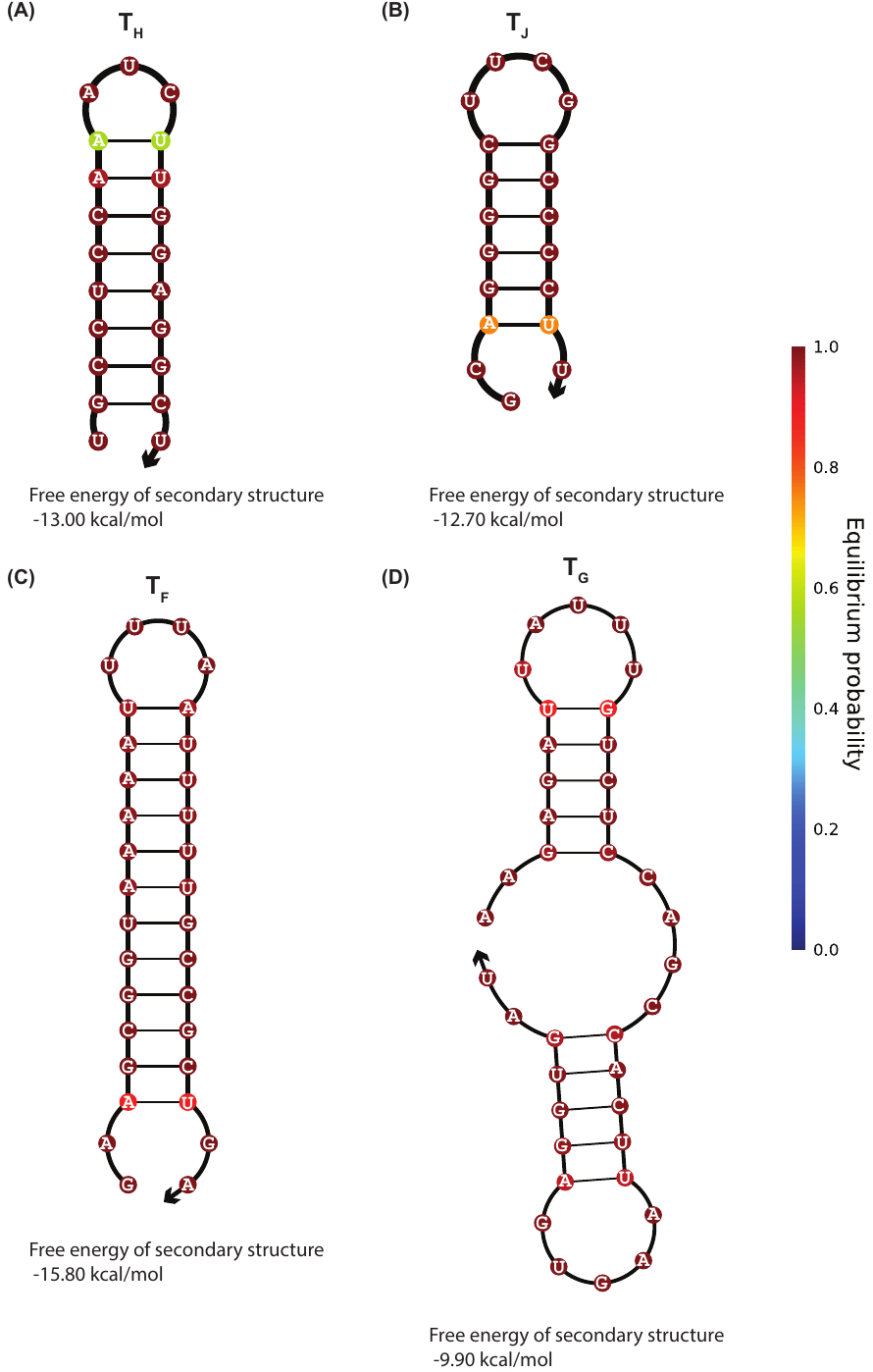

**Figure S3. Predicted structures of transcriptional terminators within øX174 genome.** (A) T_H_ terminator, (B) T_J_ terminator, (C) T_F_ terminator, (D) T_G_ terminator. Minimal free energy RNA structures (37°C) and the probability of each base pairing given by color scale, calculated by NUPACK (Zadeh et al., 2011).

**Figure S4. Computationally predicted φX174 genetic elements regulating sense transcription.** (A) Promoter prediction webtools BPROM and bTSSFINDER results (Salamov and Solovyevand, 2011). (B) Terminator prediction results from FindTerm, iTerm-PseKNC, and RibEx (Abreu-Goodger and Merino, 2005; Feng et al., 2018; Salamov and Solovyevand, 2011).

**Figure S5. Computationally predicted Rho-Utilisation (RUT) sites.** RUT sites were predicted via the RhoTermPredict (Di Salvo et al., 2019) algorithm and their sequences aligned to the φX174 genome (Genbank No. NC_001422.1).

**Figure S6. Computationally predicted φX174 genetic elements regulating antisense transcription.** (A) Promoter prediction webtools BPROM results. (B) Terminator prediction results from FindTerm, iTerm-PseKNC, and RibEx.

**Table S1.** Previously predicted sequences of φX174 promoters.

| Promoter | Sequence^a^ | Source |
| --- | --- | --- |
| *pA* | CCGTCAGGATTGACACCCTCCCAATTGTATGTTTTCATGCCTCC | (Sanger et al., 1977) |
| *pA* | AAGATTGACACCCTCCCAATTGTATGATTTCATGCCTCCA | (Sorensen et al., 1998) |
| *pB1* | TTCCTACAGGTAGCGTTGACCCTAATTTTGGTCGTCGGGTACGCA | (Sanger et al., 1977) |
| *pB2* | TTAAATAGCTTGCAAAATACGTGGCCTTATGGTTACAGTATGCCC | (Sanger et al., 1977) |
| *pB* | CCTGTTGATGCTAAAGGTGAGCCGCTTAAAGCTACCA | (Zhao et al., 2012) |
| *pB* | ATAGCTTGCAAAATACGTGGCCTTATGGTTACAGTATGCCC | (Sorensen et al., 1998) |
| *pD* | TCTCTTGTTGACATTTTAAAAGAGCGTGGATTACTATCTGAGTCC | (Sanger et al., 1977) |
| *p5211* | TGAGGTTGACTTAGTTCATCAGCAAACGCAGAATCAGCG | (Sorensen et al., 1998) |

^a^ Predicted -35 and -10 regulatory elements underlined.

**Table S2.** Stranded read mapping against NC_001422.1 reference sequence.

| **Sample** | **Read** | **Mapping Direction** | **Number of Reads** | **Proportion of Total Reads in Sample (%)** |
| --- | --- | --- | --- | --- |
| 1 | 1 | FOR | 14,850 | 0.38% |
|  |  | REV | 3,879,785 | 99.62% |
|  | 2 | FOR | 3,586,555 | 99.62% |
|  |  | REV | 13,753 | 0.38% |
| 2 | 1 | FOR | 6,830 | 0.21% |
|  |  | REV | 3,255,368 | 99.79% |
|  | 2 | FOR | 3,083,661 | 99.79% |
|  |  | REV | 6,591 | 0.21% |
| 3 | 1 | FOR | 8,968 | 0.30% |
|  |  | REV | 2,945,057 | 99.70% |
|  | 2 | FOR | 2,801,043 | 99.70% |
|  |  | REV | 8,559 | 0.30% |
